# High phenotypic and genotypic plasticity among strains of the mushroom-forming fungus *Schizophyllum commune*

**DOI:** 10.1101/2024.02.21.581338

**Authors:** Ioana M. Marian, Ivan D. Valdes, Richard D. Hayes, Kurt LaButti, Kecia Duffy, Mansi Chovatia, Jenifer Johnson, Vivian Ng, Luis G. Lugones, Han A. B. Wösten, Igor V. Grigoriev, Robin A. Ohm

**Affiliations:** Microbiology, Department of Biology, Faculty of Science, Utrecht University, Padualaan 8, 3584 CH, Utrecht, The Netherlands; U.S. Department of Energy Joint Genome Institute, Lawrence Berkeley National Laboratory, Berkeley, CA 94720, USA

## Abstract

*Schizophyllum commune* is a mushroom-forming fungus notable for its distinctive fruiting bodies with split gills. It is used as a model organism to study mushroom development, lignocellulose degradation and mating type loci. It is a hypervariable species with considerable genetic and phenotypic diversity between the strains. In this study, we systematically phenotyped 16 dikaryotic strains for aspects of mushroom development and 18 monokaryotic strains for lignocellulose degradation. There was considerable heterogeneity among the strains regarding these phenotypes. The majority of the strains developed mushrooms with varying morphologies, although some strains only grew vegetatively under the tested conditions. Growth on various carbon sources showed strain-specific profiles. The genomes of seven monokaryotic strains were sequenced and analyzed together with six previously published genome sequences. Moreover, the related species *Schizophyllum fasciatum* was sequenced. Although there was considerable genetic variation between the genome assemblies, the genes related to mushroom formation and lignocellulose degradation were well conserved. These sequenced genomes will provide a solid basis for functional genomics analyses of the strains of *S. commune*.

## Introduction

*Schizophyllum commune* is a lignocellulose-degrading and mushroom-forming fungus in the class Agaricomycetes, phylum Basidiomycota. It has a wide geographic distribution and has been found on all continents. It is usually found on dead wood, and although it has been described as a white rot fungus (i.e., it degrades all components of wood), it has more recently been classified as uncertain wood decay type (UWD) (Floudas et al., 2020). *S. commune* is used as a model organism to study mushroom formation, mating type loci, and lignocellulose degradation. It completes its life cycle in ten days, it has the ability to form mushrooms on defined synthetic media, the 38.7 Mb genome has been sequenced (Marian et al., 2022; Ohm et al., 2010) and several molecular tools are available, including efficient CRISPR/Cas9 genome editing (De Jong et al., 2010; Vonk et al., 2019).

The first sequenced strain was H4-8 (Ohm et al., 2010), which is the reference strain used in our studies. More recently, additional strains were sequenced (Baranova et al., 2015; Boiko, 2022; Marian et al., 2022; Seplyarskiy et al., 2014). These analyses have shown that *S. commune* is a very polymorphic species, both phenotypically and genotypically. Still, only a limited number of *S. commune* strains have been sequenced and annotated to date. Therefore, sequencing additional strains is required to gain more insights into the phylogeny and functional genomics of the species. Specifically, additional genomes of strains with diverse phenotypes regarding mushroom development and lignocellulose degradation may allow us to identify the genetic differences that underlie the phenotypic diversity.

Mushrooms are the reproductive structures of *S. commune* and many other members of the class Agaricomycetes. The decision to form mushrooms is influenced by environmental stimuli, including light intensity, CO_2_ concentration, temperature, and humidity. However, the effect of these stimuli is species-specific. In the case of *S. commune*, high concentrations of CO_2_ have been described as an inhibitor of fruiting body formation (Niederpruem, 1963; Raudaskoski and Viitanen, 1982), while blue light induces fruiting body formation (Ohm et al., 2013).

Several transcription factors are known to regulate mushroom formation. For example, a blind strain unable to fruit is obtained if either one or both of the genes of the blue light receptor White Collar Complex (WC-1 and WC-2) is inactivated (Ohm et al., 2013). Other transcription factors involved in aspects of mushroom development in *S. commune* are Hom1, Hom2, Fst1, Fst3, Fst4, Bril, C2h2, Gat1, Zfc7 and Tea1 (Ohm et al., 2011, 2010; Pelkmans et al., 2017; Vonk and Ohm, 2021, 2018). However, there are over 200 putative transcription factors whose function remains largely unknown. Few classes of proteins are known to play a structural role in mushroom development, although hydrophobins are a notable exception. They are fungal-specific, relatively short, and hydrophobic proteins that facilitate aerial growth and mushroom development (Wösten, 2001). Several hydrophobin genes are highly expressed during fruiting body formation in *S. commune* (Krizsán et al., 2019; Ohm et al., 2010). Understanding the roles of these genes in mushroom development may allow for more efficient cultivation of mushrooms. Moreover, little is known about differences in mushroom development between strains, and the genes responsible for these developmental phenotypes.

Members of the class Agaricomycetes are particularly good at degrading lignocellulose and their enzymatic potential has been studied extensively (Ohm et al., 2014; Riley et al., 2014). In nature, *S. commune* grows on wood and its genome encodes a range of carbohydrate-active enzymes (CAZymes). Wood mainly consists of the polymers cellulose, hemicellulose, pectin and lignin. CAZymes have the potential to degrade all these components and are classified into families of Glycoside Hydrolases (GHs), Glycosyl Transferases (GTs), Polysaccharide Lyases (PLs), Carbohydrate Esterases (CEs), and Auxiliary Activities (AAs) (Drula et al., 2022). The conserved transcription factor Roc1 regulates the expression of genes encoding cellulase-degrading enzymes (Marian et al., 2022).

Here, we studied the diversity between 16 dikaryotic and 18 monokaryotic strains by systematically phenotyping aspects of mushroom development, lignocellulose degradation and interaction profile. Moreover, we sequenced the genomes of seven monokaryotic strains of *S. commune* and one strain of the related species *S. fasciatum*. We performed a comparative genomics analysis with the genome sequences of eight previously published strains.

## Results

### The genome assemblies of *S. commune* strains reveal high sequence diversity

The genotypic diversity of the genus *Schizophyllum* was assessed by sequencing seven monokaryotic (i.e., haploid) strains of *S. commune* and one monokaryotic strain of the related species *S. fasciatum*. These were all obtained by protoplasting dikaryotic (i.e., heterokaryotic) strains collected from various locations in the world to ensure a degree of geographic diversity (Table 1). Three previously published sequenced strains were also included in the analysis. The statistics of the assemblies, gene predictions and functional annotations are listed in Table S1.

**Table 1.**
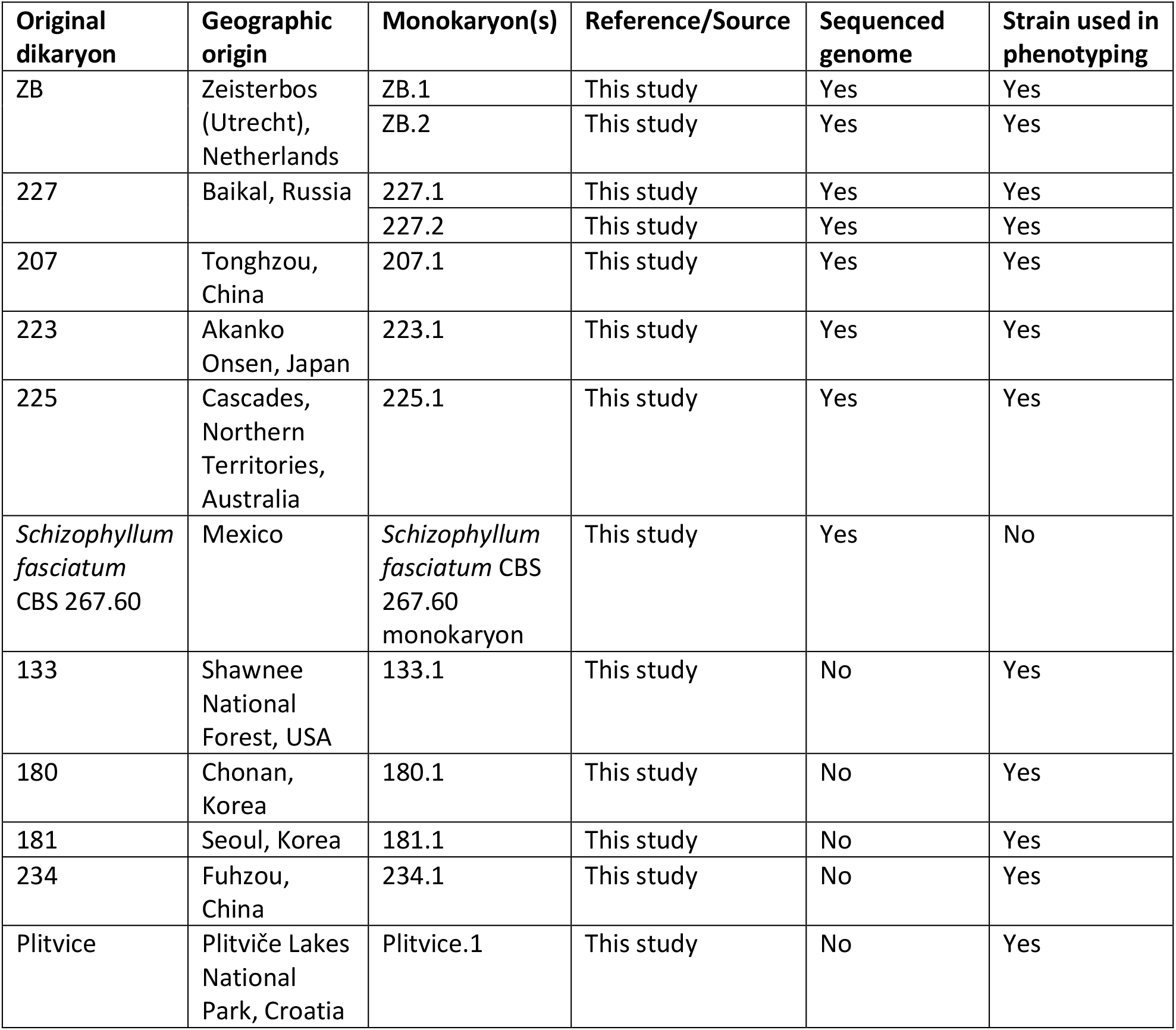
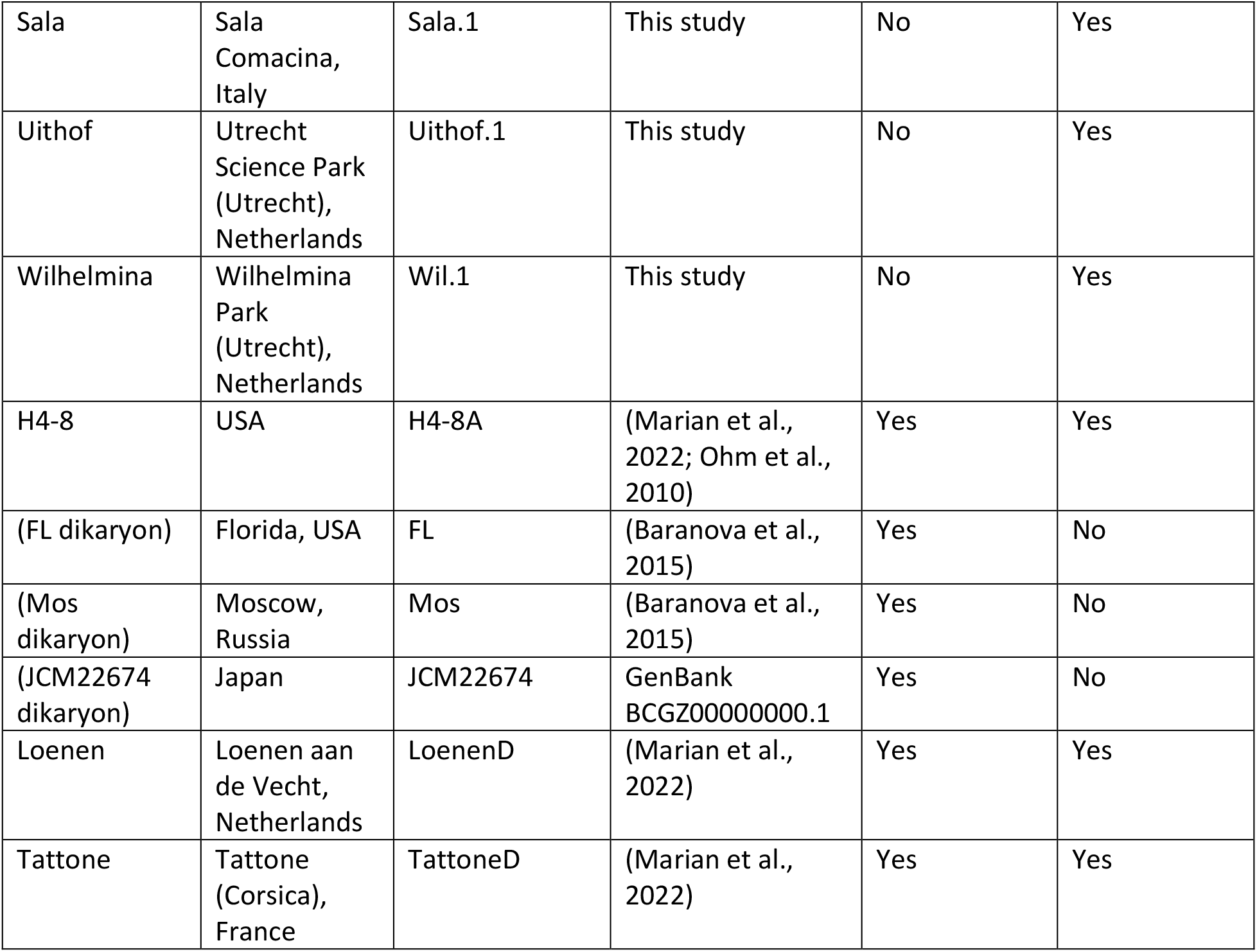
Dikaryotic strains and their corresponding monokaryons.

The size of the assembly of *S. fasciatum* is considerably smaller (30.4 Mbp) than those of *S. commune* (38.7 Mbp for reference strain H4-8). It also encodes fewer genes (11722 and 16204 genes for *S. fasciatum* and *S. commune* H4-8, respectively). Nevertheless, the BUSCO completeness score indicates that the genome is complete (albeit rather fragmented). Little is known about the lifestyle of this close relative of the well-studied *S. commune*.

A phylogenetic tree was reconstructed, with the related species *Auriculariopsis ampla* and *Fistulina hepatica* as outgroups (Figure 1A). *S. fasciatum* is a sister taxon to the clade of *S. commune* strains. Generally, the *S. commune* strains cluster by geographical distribution. The European and North-American strains are both monophyletic, but the Asian strains are paraphyletic. The single Australian strain is most closely related to an Asian (more specifically, Japanese) strain. Even though monokaryons ZB.1 and ZB.2 both originate from the dikaryon ZB, they are not closely related. Similarly, 227.1 and 227.2 also do not cluster closely together.

**Figure 1.**
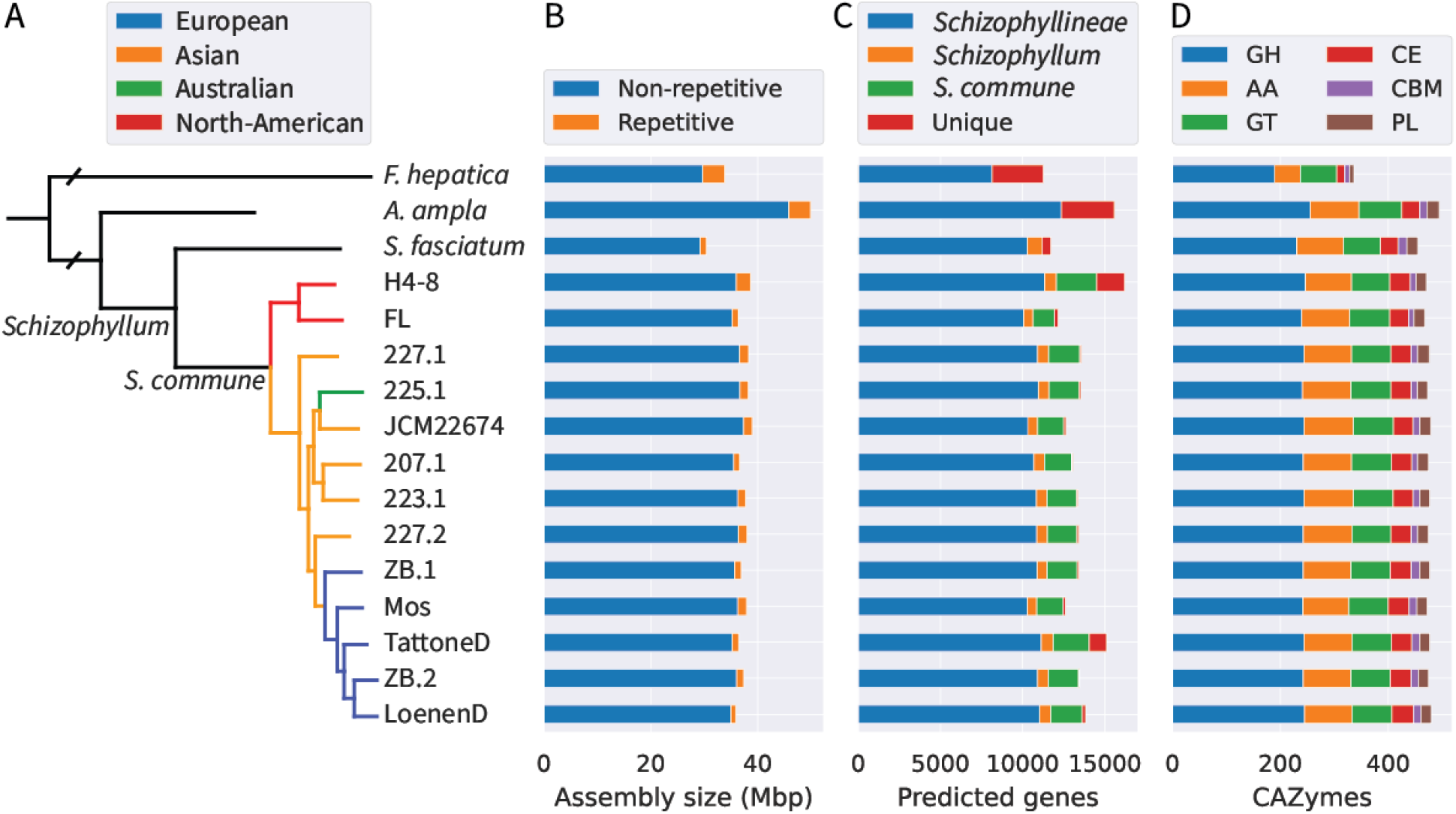
**A**. Phylogenetic tree of the genus *Schizophyllum. S. fasciatum* is a sister taxon to the clade of *S. commune* strains, which cluster together largely by their geographic origin. The species *Fistulina hepatica* and *Auriculariopsis ampla* are used as outgroup. The tree is rooted on *F. hepatica* and this branch is not drawn to scale. **B**. The assemblies of the *S. commune* strains have similar sizes, but the assembly of *S. fasciatum* is considerably smaller. **C**. The number of predicated genes is similar for the *S. commune* strains, with higher counts for strains H4-8 and TattoneD, which can be explained by a difference in method of gene prediction. The genome of *S. fasciatum* encodes considerably fewer genes. **D**. The number of predicted carbohydrate-active enzymes (CAZymes) is remarkably similar for all *S. commune* strains.

The assemblies of the *S. commune* strains were similar in size (Figure 1B). An all-versus-all comparison shows that there is considerable variation on assembly-level between the strains (Figure 2A). For example, there is less than 80% sequence identity between the assemblies of H4-8 and the European strains. To identify conserved regions in the assemblies, we compared all *S. commune* strains to the reference strain H4-8 and calculated the average conservation across all strains (Figure 2B). There is considerable variation and especially the putative sub-telomeric regions at the ends of the scaffolds tend to be less conserved. Nevertheless, the region on the left arm of scaffold 2 is relatively well conserved in most strains (Figure S1). Interestingly, this region contains the mating type A (matA) locus.

**Figure 2.**
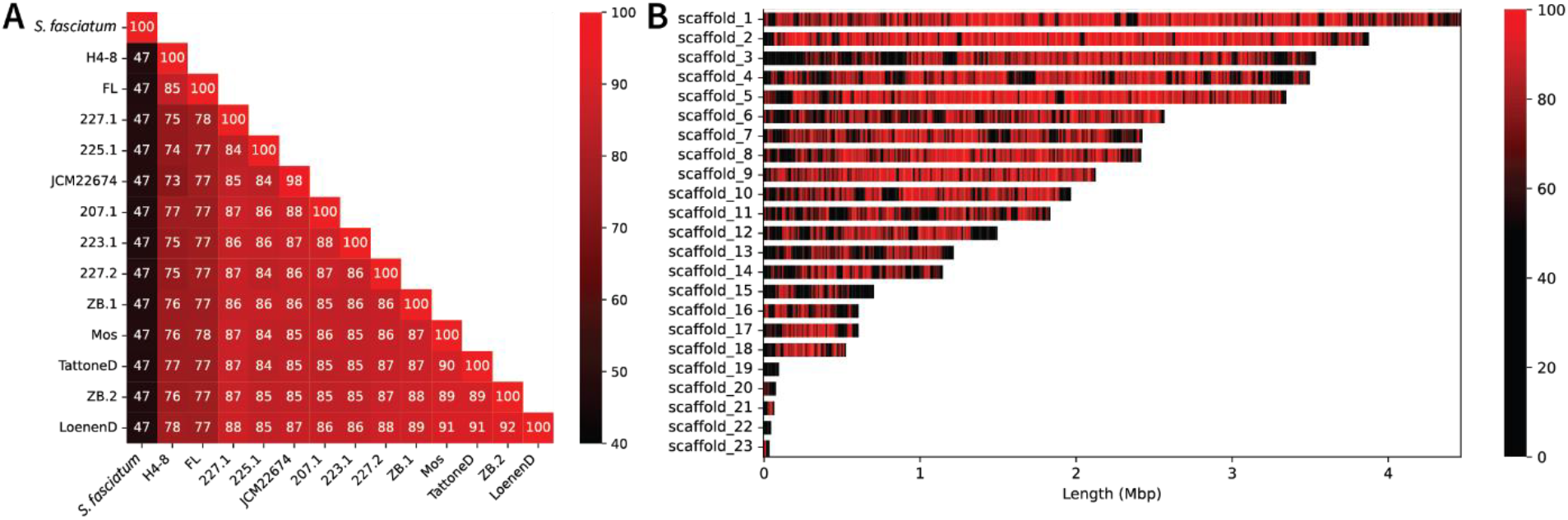
**A**. Conservation between the assemblies of the strains in the genus *Schizophyllum*. The conservation is expressed as percentage of sequence identity. Even though the *S. commune* strains are of the same species, their assemblies display a high degree of variation. **B**. The average conservation between the reference assembly of strain H4-8 and the assemblies of the other 12 *S. commune* strains. The scaffolds of the reference assembly are visualized along the y-axis, and the scaffold lengths are indicated on the x-axis. The individual pair-wise comparisons of the assemblies are visualized in Figure S1.

The gene counts vary between the *S. commune* strains (Figure 1C), although it is likely that this is largely due to the annotation method. Strains H4-8 and TattoneD were annotated using RNA-Seq data to aid gene prediction, while this was not the case for the other strains. Expression data is known to identify additional genes that may otherwise be missed (Grigoriev et al., 2006). The strains that were annotated with the JGI annotation pipeline show similar gene counts, while strain FL, Mos and JCM22674 were annotated separately. Most genes are conserved across the species in the analysis, although there are also genes conserved only in the genus *Schizophyllum* or the species *S. commune* (Figure 1C). Relatively few genes are unique to a strain, with the exception of strain H4-8 and TattoneD, which were annotated using expression data and therefore likely include genes that were missed in other strains. Overall, the gene counts are rather similar between the strains.

The putative mating type genes were identified based on their functional annotations (Table S2). As expected, these mating type genes are organized in mating type loci. However, due to the relatively high number of scaffolds in the Illumina-sequenced assemblies, the mating type loci are generally fragmented. Despite the smaller assembly size and lower gene count of *S. fasciatum*, the number of pheromones and pheromone receptors is considerably higher than for the strains of *S. commune*.

### Fruiting body development shows high phenotypic diversity between strains

The dikaryotic reference strain H4-8 develops mushrooms when grown for 12 days from a point inoculum on solid SCMM, at 25°C, in a 16 h light / 8 h dark cycle. The dikaryotic wild isolate strains were screened under the same conditions and their mushroom development phenotype was analyzed (Figure 3).

**Figure 3.**
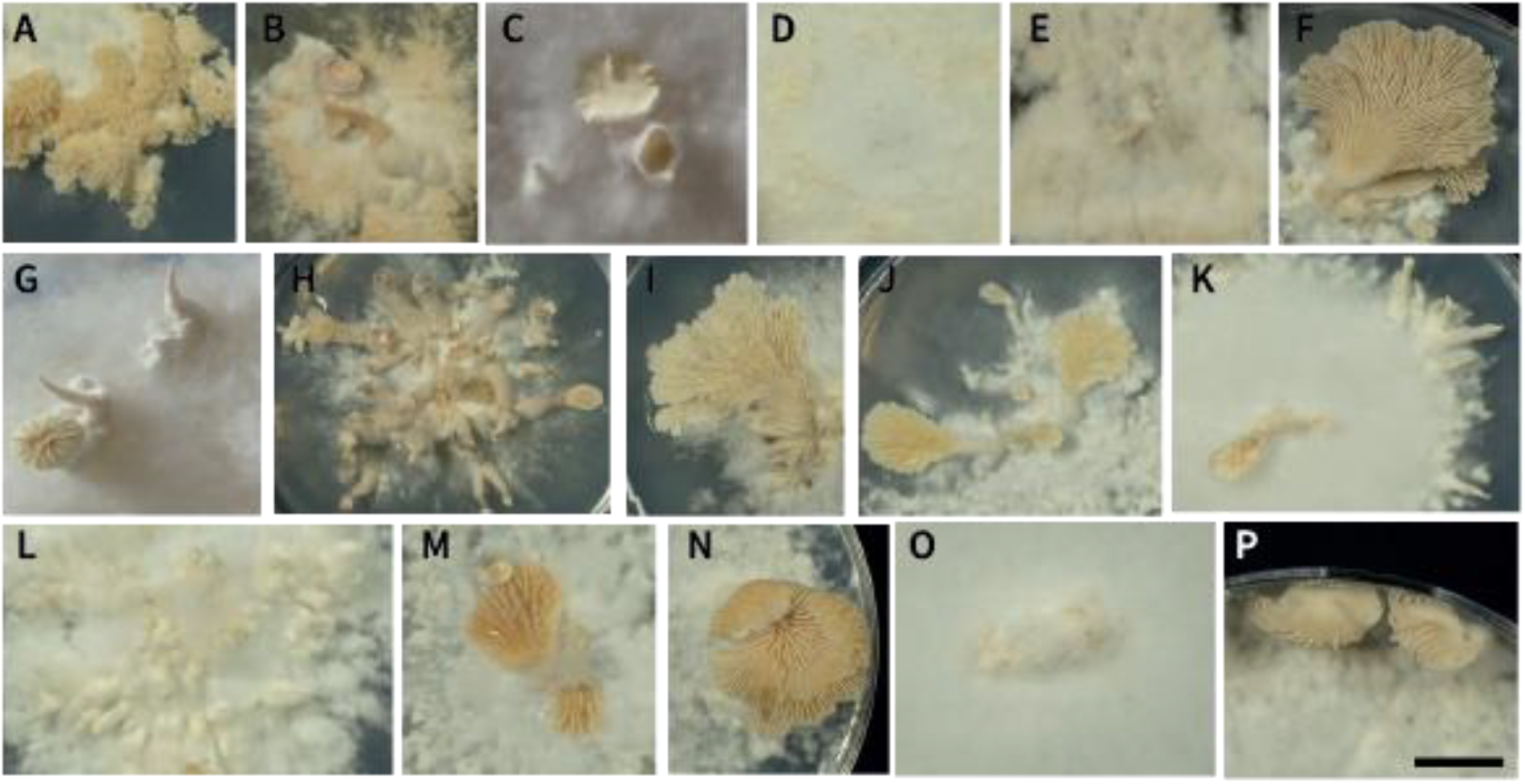
There is considerable diversity in fruiting body morphology among dikaryotic strains of S. commune. **A**. H4-8; **B**. Tattone; **C**. Loenen; **D**. ZB; **E**. 227; **F**. 207; **G**. 223; **H**. 225; **I**. 133; **J**. 180; **K**. 181; **L**. 234; **M**. Plitvice; **N**. Sala; **O**. Uithof; **P**. Wilhelmina. The strains were grown at 25°C in a 16 h light / 8 h dark cycle. The pictures were taken after 12 days of growth. The scale bar in (P) represents 1 cm.

Mushroom formation varied considerably between the strains. Reference strain H4-8 (Figure 3A) formed an abundance of small cup-shaped and stipeless fruiting bodies growing in a ring around the center. Most of the wild isolate strains also fructified in these conditions, but there were two strains that did not go further than the mycelium/aggregates stage (ZB and Uithof). The strains that formed mushrooms displayed a range of shapes, sizes, abundance and colors. Loenen and Tattone both exhibited tube-shaped fruiting bodies. Loenen (Figure 3C) formed a considerably larger amount of vegetative mycelium than Tattone (Figure 3B). In strain 227 (Figure 3E), fruiting body development was arrested at the aggregates stage and after initiation of fruiting body formation a dark brown fungal pigment was produced in the growth medium. Strain 223 (Figure 3G) developed fruiting bodies in the center of the plate and formed either stipes or mature mushrooms. Strain 225 (Figure 3H) exhibited a multitude of fruiting bodies originating in the center of the colony, with long stipes and few mushrooms that fully opened. Strain 234 (Figure 3L) displayed a multitude of aggregates and stipes that did not develop further to the mushroom stage. Strains 133 (Figure 3I), 207 (Figure 3F) and Sala (Figure 3N) developed similar large fan-shaped fruiting bodies attached to a short stipe. Similarly, strains 180 (Figure 3J) and Plitvice (Figure 3M) also developed fan-shaped mushrooms concentrated in the center of the plate, although smaller than in strains 133, 207 and Sala. Strain 181 (Figure 3K) predominantly formed mushrooms at the periphery of the colony, which is not generally seen in any other strain. Strain Wilhelmina (Figure 3P) produced unusually shaped fruiting bodies at the periphery of the colony.

### The effect of CO2 on fruiting body development varies between the strains

The CO_2_concentration plays an important role in developmental decisions in mushroom-forming fungi. In the reference strain H4-8, fruiting bodies generally develop in low (i.e. 400 ppm) CO_2_concentration, whereas their formation is inhibited at high (5%) CO_2_ concentrations. The dikaryotic wild isolate strains were screened under the same conditions and their mushroom development phenotype was analyzed (Figure 4). At a high CO_2_ concentration, the majority of strains did not develop mushrooms but instead only formed vegetative mycelium. In contrast, strains 207, 225, 180 and 234 did produce fruiting bodies at a high CO_2_ concentration. However, mushroom shape was considerably different: the stipes were longer, the cap was smaller and less spore-forming tissue was developed, compared to when grown at low CO_2_ concentration. Especially for strain 225 the difference between low and high CO_2_concentration regarding colony and mushroom morphology was relatively small.

**Figure 4.**
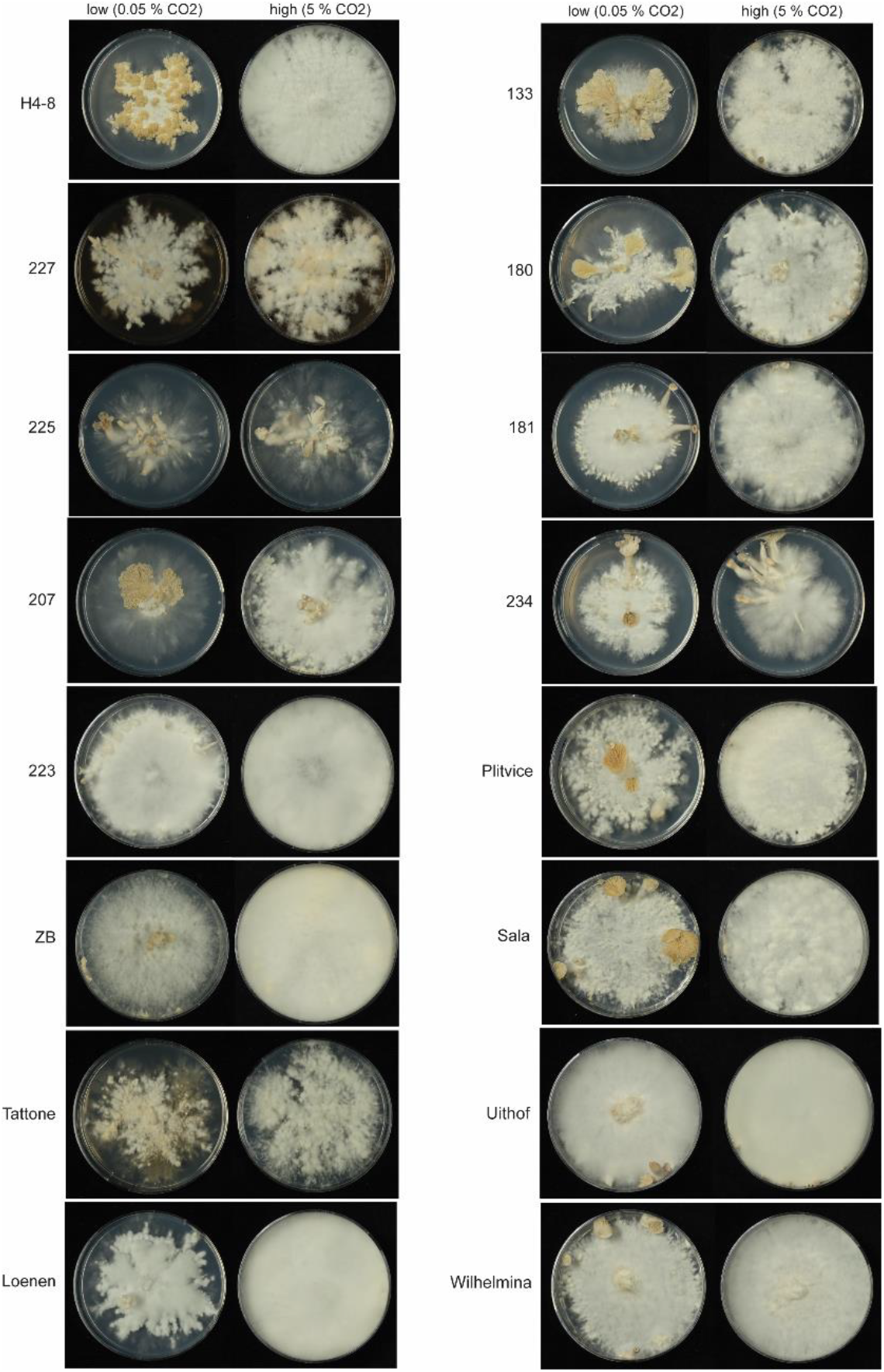
Morphological comparison of *S. commune* dikaryotic strains under different CO_2_ concentrations. The strains were grown on for 12 days at 25°C under continuous light.

**Figure 5.**
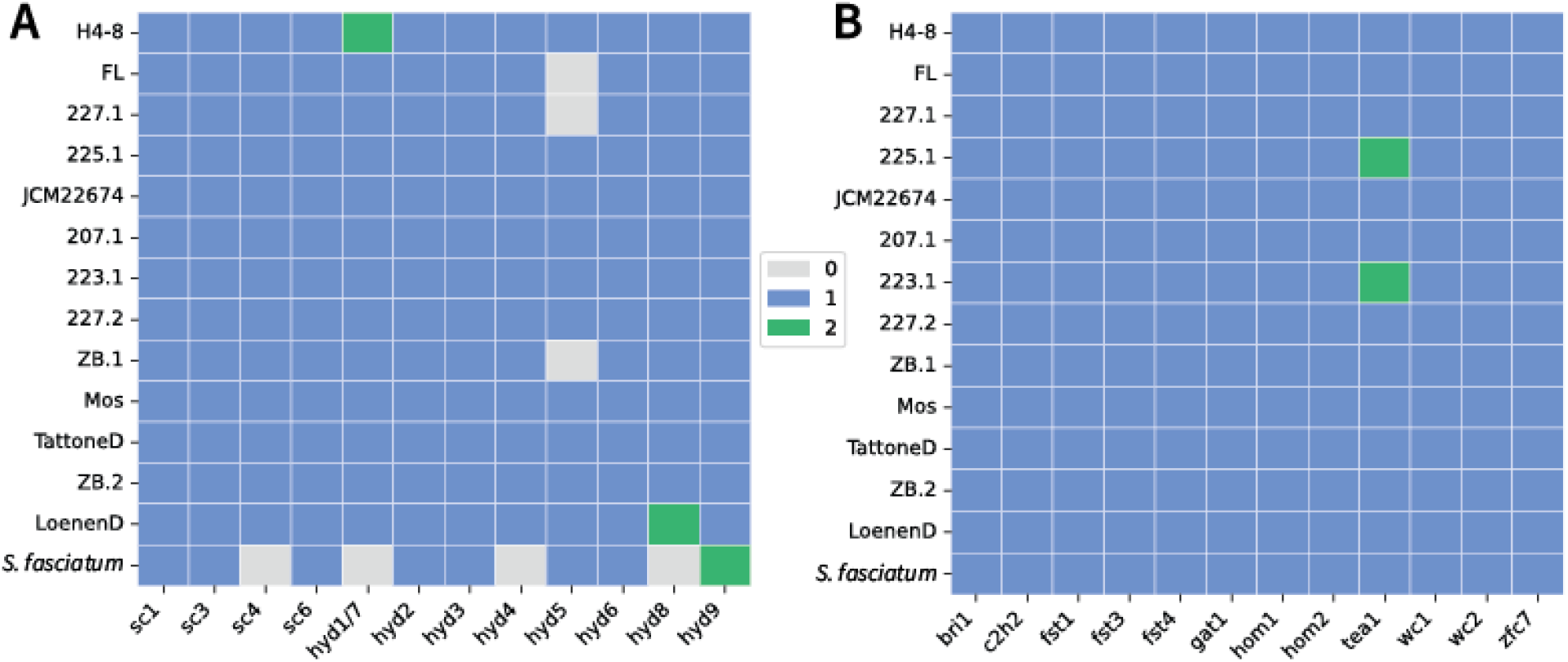
**A**. Diagram depicting the number of hydrophobin orthologs in the genus *Schizophyllum*. **B**. The number of orthologs of regulators with a role in mushroom formation in the genus *Schizophyllum*.

### Hydrophobins and regulators involved in mushroom development

The wide variation in mushroom development phenotypes likely has a genetic basis. Therefore, we analyzed the conservation of known genes and gene families (hydrophobins and transcription factors) involved in mushroom development across the strains of *S. commune* and in *S. fasciatum*.

The genome of reference strain H4-8 encodes 13 hydrophobin genes (*sc1, sc3, sc4, sc6, hyd1-9*). The hydrophobins of the newly sequenced strains were compared to these and their conservation was determined. Hydrophobin genes are generally conserved in the various strains of *S. commune*, with some notable exceptions. Genes *hyd1* and *hyd7* of the reference strain H4-8 were in fact the result of a recent duplication that only took place in this strain, and we therefore combined these into group *hyd1/7*. In strain LoenenD, gene *hyd8* has also undergone a recent duplication. Gene *hyd5* is lost in the strains FL, 227.1 and ZB.1. The genome of *S. fasciatum* encodes fewer hydrophobins and for example lacks an ortholog of *sc4*, which is involved in mushroom development. However, *hyd9* is duplicated in this species.

Transcription factors involved in the regulation of mushroom development are generally well conserved in the various strains of *S. commune*. A notable exception is *tea1* which has duplicated in strain 223.1 and 225.1. All these regulators are also conserved in *S. fasciatum*.

### Growth on wood-associated carbon sources varies between wild isolate strains

*S. commune* has the ability to utilize various carbon sources. We studied the growth on nine carbon sources that were associated with wood and its components: beech wood, cellulose (a component of wood), cellobiose (a breakdown product of cellulose), glucose (a sugar that is easily metabolized, and a breakdown product of cellulose), pectin (a component of wood), xylan (a type of hemicellulose and a component of wood), xylose (a breakdown product of xylan), and maltose and starch (both are not associated with wood, but are relatively easily metabolized).

Reference strain H4-8 grew on beech wood, although to a limited extent. It displayed considerably slower growth than strains 227.1, 223.1, Sala.1 and 181.1, which formed a denser mycelium. On the other hand, strains ZB1, ZB2 and 227.2 grew poorly on wood, although a sparse mycelium was visible throughout the substrate.

On cellulose, the strains displayed relatively sparse growth compared to other carbon sources (except beech wood). However, reference strain H4-8, 207.1, 227.1, 223.1 and 181.1 grew better than the other strains. Generally, these strains also grew better on wood than most of the other strains, although there is a notable exception (Sala.1). Growth on cellobiose was more similar between the strains, although strain 227.1 displayed considerably slower growth resulting in a smaller colony. Xylan (a type of hemicellulose) and its breakdown product xylose were readily used by most strains. Notable exceptions include strains LoenenD and 227.1, which both displayed sparser growth on these two carbon sources. Growth on pectin was generally less fast than on the other carbon sources, and there was little difference between the strains. However, strain ZB.2 grew considerably faster than H4-8.

Glucose, maltose and starch are carbon sources that are considerably easier to metabolize than the predominantly polymeric carbon sources described above. The various strains displayed little variation during growth on these carbon sources.

### Correlation between growth on cellulose and cellulase activity

We determined whether the growth profile of the strains on solid medium with cellulose as sole carbon source could be explained by a difference in cellulase enzyme activity. Since the enzyme activity is difficult to measure in solid medium, we performed these experiments in liquid shaken cultures. After 6 days of growth the supernatant was collected, and the total cellulase enzyme activity was measured using the filter paper activity assay. In general, there was a high variation between cellulase enzyme activity of the strains (Figure 7). However, high cellulase activity often correlated with growth on solid cellulose and wood medium. Reference strain H4-8, and strains 223.1, 207.1, 180.1 and 181.1 had a relatively high cellulase activity in the growth medium and a relatively strong growth on solid cellulose medium (Figures 6 and 7). This shows that for these strains the strong growth on cellulose may (at least in part) be explained by the increased production of cellulase enzymes. However, strain 225.1 also showed a relatively high cellulase activity, but displayed almost no growth on solid cellulose medium (although growth on wood was relatively good). Inversely, strain 234.1 showed very little cellulase activity in liquid medium, but growth on solid cellulose was relatively good. It should be noted that growth on wood was very poor for strain 234.1. Possibly, these discrepancies may be explained by the differences between growth in liquid and on solid medium.

**Figure 6.**
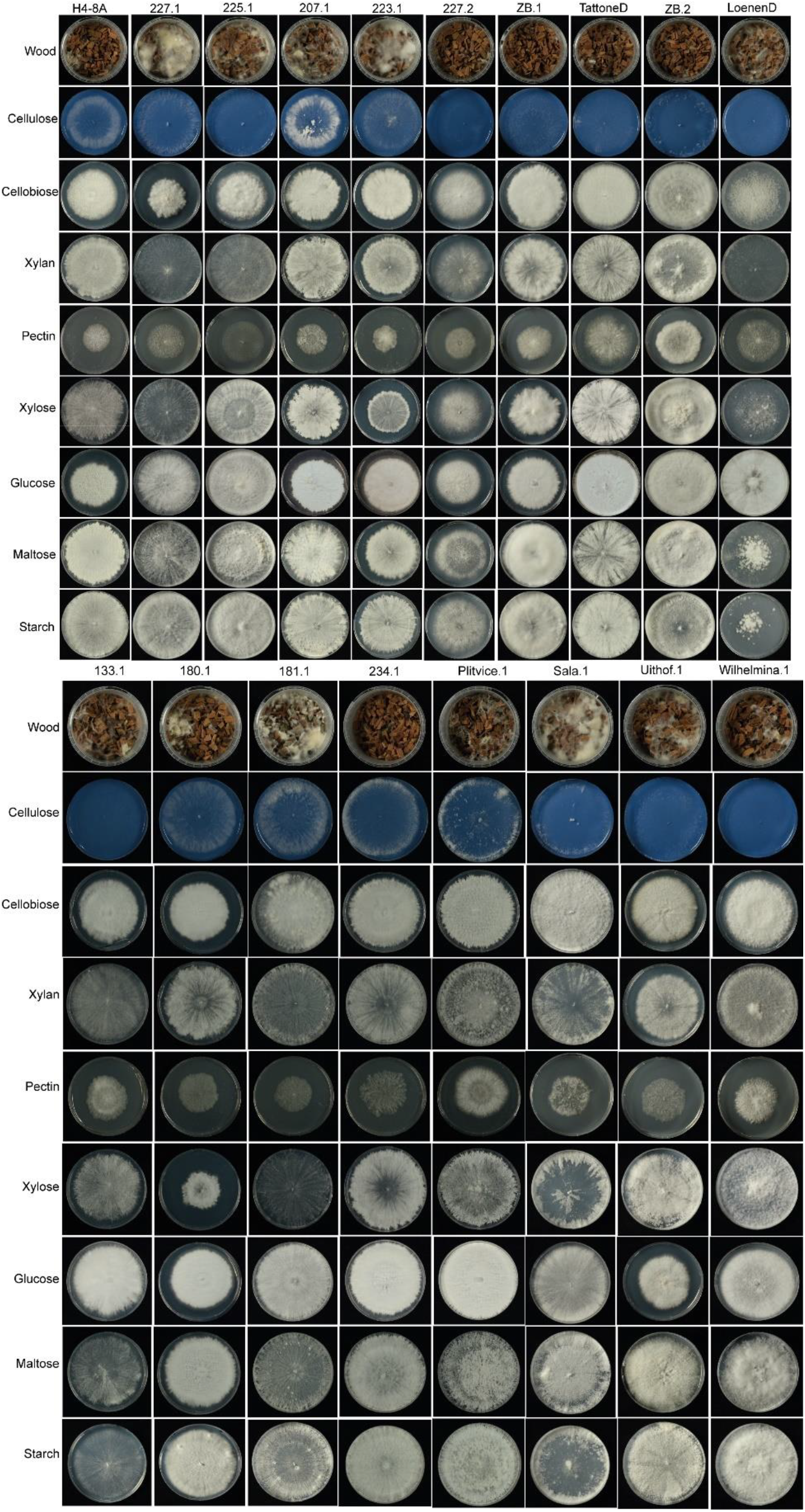
Growth analysis of *S. commune* monokaryotic strains on nine different carbon sources. Pictures were taken after seven days (or 32 days in the case of wood) of growth at 30°C.

**Figure 7.**
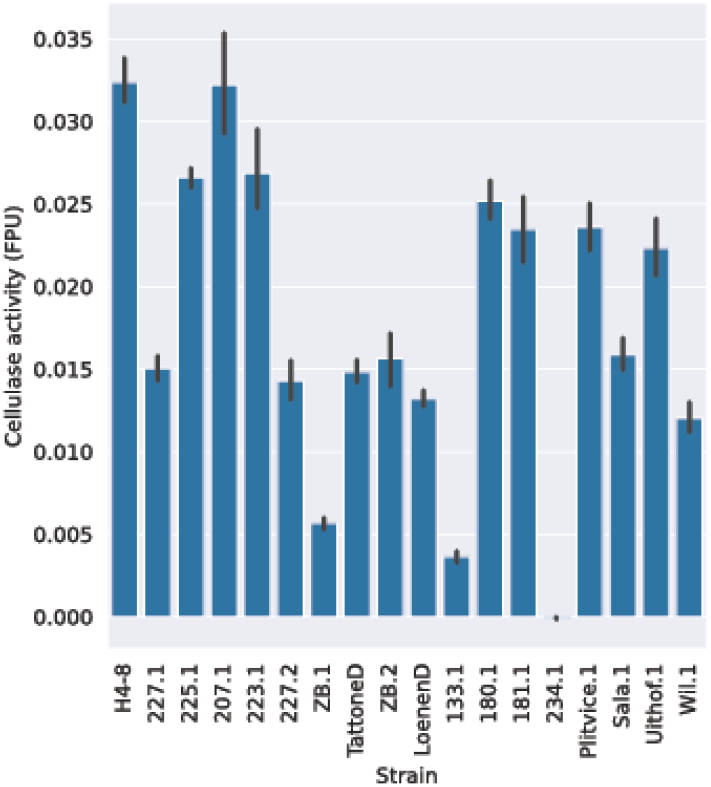
Total cellulase activity during growth in liquid culture with cellulose as sole carbon source. There is considerable heterogeneity between the strains of *S. commune*.

### There is little variation in CAZyme count despite differences in growth profile and cellulase activity

We determined whether we could identify an underlying genetic component that may explain the different growth profiles on the various wood-related carbon sources, as well as the different cellulase activities of the strains. Specifically, we determined the number of genes encoding CAZymes in the genome assemblies of the strains.

The genomes of the sequenced *S. commune* strains encode a similar number of putative CAZymes (Figure 1D, Table S3), ranging from 481 in reference strain H4-8 to 495 genes in strain JCM22674. These differences are relatively small, even if we take into account the genome-size variance. All strains encode glycoside hydrolases (GHs), glycosyl transferases (GTs), polysaccharide lyases (PLs), carbohydrate esterases (CEs), auxiliary activities (AAs) and carbohydrate-binding module (CBM) families. The GH and AA families are the most represented among the CAZymes and these are primarily (but not exclusively) associated with degradation of wood, cellulose, hemicellulose, pectin, lignin and starch.

These relatively small differences in CAZyme gene counts cannot explain the relatively large differences in growth profile and cellulase activities observed between some of the strains. It is therefore likely that the observed differences are caused at the regulatory level or the activity of individual cellulase enzymes. The regulator of cellulose degradation Roc1 is conserved in all strains, as well as in *S. fasciatum*.

## Discussion

There is considerable genomic and phenotypic diversity among the *S. commune* strains. The genome sequences of the 7 newly sequenced strains and the 6 previously published strains showed a high degree of genetic diversity. This is also reflected in the high degree of phenotypic diversity in various aspects of mushroom development and lignocellulose degradation.

*S. commune* was previously described as among the most polymorphic of all studied eukaryotic species (Baranova et al., 2015) and the newly sequenced strains also show considerable genomic diversity. The phylogenetic analysis showed that genetic distance largely correlates with geographic distance. Strains from the Eastern and Western hemispheres form distinct clades. Moreover, it appears that *S. commune* spread from Asia into Europe and Australia. This largely confirms population analyses performed on a smaller set of alleles (James et al., 2001), and showed that the oceans are likely barriers to dispersal of this species (James et al., 1999). Sequencing additional strains from geographically distinct areas, as well as including additional previously published strains (Baranova et al., 2015; Kim et al., 2021; Liu et al., 2022; Seplyarskiy et al., 2014) (Baranova et al., 2015; Kim et al., 2020; Liu et al., 2022; Seplyarskiy et al., 2014), will yield more insights into the population dynamics of *S. commune*.

It is remarkable that neither ZB.1 and ZB.2, nor 227.1 and 227.2 cluster closely together, given that each of these pairs are monokaryons derived from one dikaryon by protoplasting. Apparently, these dikaryons originated from monokaryons that were not closely related and possibly travelled a considerable distance. These two examples show that outcrossing happens frequently in *S. commune*, although sequencing additional dikaryons is required to further determine the extent. This large genetic diversity between the genomes of the two nuclei cannot be resolved by Illumina short-read sequencing, as we have been unsuccessful at sequencing and assembling dikaryotic genomes using this approach (results not shown). Long-read sequencing techniques (e.g., PacBio or Oxford Nanopore) will likely be able to resolve these issues.

*S. fasciatum* is the first other species in the genus *Schizophyllum* (besides *S. commune*) that has been sequenced. Little is known about this strain, except that is has been collected from Mexico. It is not known whether the high genetic diversity found in *S. commune* is restricted to only this species, or whether *S. fasciatum* and other species show a similar pattern.

There was considerable heterogeneity in mushroom development phenotypes between the wild isolate dikaryons. These wild isolates have been collected from nature as (apparently) recognizable mushrooms of the species *S. commune*. Nevertheless, some strains did not develop mushrooms when grown in laboratory conditions. However, the conditions that were used in the laboratory (i.e., growth in Petri dishes, glucose as carbon source instead of wood, absence of competitors, etc) were considerably different from nature. Moreover, since the strains were collected in different geographic locations, the strains have likely adapted to their local environments (e.g., temperature, humidity, day/night cycle, light intensity, etc) and wood substrates, which may have resulted in considerable heterogeneity between the strains. These adaptations have likely resulted from genetic differences. Several gene families have been previously associated with mushroom development in *S. commune*. Hydrophobins play an important role in allowing the hyphae to grow up into the air, which is a first step in mushroom development. Moreover, they coat the air channels in the mushrooms, preventing those from filling up with water (Wösten, 2001). Several transcription factors have been shown to regulate various aspects of mushroom development. For this reason, we analyzed the conservation of both hydrophobins and specific transcription factors in the newly sequenced strains. Remarkably, despite the high genetic diversity, there was little difference in gene counts of hydrophobins and orthologs of known regulators of mushroom development. Therefore, we were unable to link the mushroom development phenotypes to specific differences in genotype. However, it should be noted that we have performed the mushroom development phenotyping with dikaryotic strains, and not all genomes of the corresponding (parental) monokaryotic strains are available (Table 1).

Similarly, we found large heterogeneity between the strains in the growth profiles of the strains on various carbon sources. Several strains grew well on most carbon sources, while other strains grew relatively poorly. Possibly, this is the result of adaptations of the strains to their niche. There was very little difference in the CAZyme gene counts in the various strains. Possibly, the heterogeneity may be explained by differences in levels of gene expression, for example caused by differences in the regulatory pathways leading to CAZyme expression. Alternatively, there may be differences in the enzyme activity of individual CAZymes. Furthermore, unknown CAZymes may play a role in lignocellulose degradation.

The phenotypic diversity between strains opens the door to using a functional genomics approach to link genotype to phenotype. Although we identified considerable differences in both the genotype and phenotype between the strains of *S. commune*, we were generally unable to link a specific genotype to a specific phenotype. The number of genetic differences on genome-level was generally too high, whereas the gene counts of specific families were generally very similar. The number of strains in our analysis is insufficient to perform a genome-wide association study (GWAS), which may be remedied by sequencing additional strains of our collection. An alternative approach is bulked segregant analysis (Wu et al., 2018), which entails crossing compatible strains with different phenotypes for a trait of interest, selecting offspring and dividing these into pools with opposing phenotypes. Sequencing these pools may reveal which loci are responsible for the observed phenotypes. Moreover, we have shown that comparative transcriptomics may be used to compare gene expression between strains growing on several carbon sources, which allowed us to identify the transcription factor Roc1 (Marian et al., 2022).

In conclusion, the genome sequences generated in this study will serve as a foundation for more in-depth analyses to identify the genetic basis of phenotypic diversity among the strains.

## Material and Methods

### Strains, media composition and culture conditions

The *S. commune* wild isolate strains used in this study were collected from various locations (Table 1). All strains were collected prior to 2014. *S. fasciatum* CBS 267.60 was obtained from the Westerdijk Fungal Biodiversity Institute and originally collected from Mexico. The corresponding monokaryotic strains (as indicated in Table 1) were obtained after protoplasting the dikaryotic mycelium using a *Trichoderma harzianum* lytic enzyme mix as previously described (Van Peer et al., 2009).

*S. commune* monokaryotic strains were grown at 30°C on a medium comprising per L: 22 g glucose monohydrate, 1.32 g (NH_4_)_2_SO_4_, 0.5 g MgSO_4_·7H_2_O, 0.12 mg thiamine, 1 g K_2_HPO_4_, 0.46 g KH_2_PO_4_, 5 mg FeCl_3_.6H_2_O, trace elements (Whitaker, 1951) with 1.5% agar. For cultures with other carbon sources, glucose was replaced with 1% (w/v) Avicel, 1% (w/v) cellobiose, 1% (w/v) xylan from corn cob, 1% (w/v) pectin from apple, 1% (w/v) starch from potato, 2.2% (w/v) xylose, 2.2% (w/v) maltose monohydrate. To improve the visualization of the colonies grown on Avicel, the media was supplemented with 20 μg μl-1 Remazol Brilliant Blue R. For growth on wood we used 60 g of rinsed beech wood chips in plastic boxes (9 cm diameter) with a lid containing a filter, and the samples were grown for 32 days.

For the cellulase enzyme activity assay, liquid cultures were grown in Erlenmeyer flasks at 30°C, shaking at 200 rpm, in a medium without Avicel (MwA) comprising 1.32 g (NH_4_)_2_SO_4_, 0.5 g MgSO_4_·7H_2_O, 0.12 mg thiamine, 1 g K_2_HPO_4_, 0.46 g KH_2_PO_4_, 5 mg FeCl_3_.6H_2_O and trace elements (Whitaker, 1951). Next, 1 g Avicel was added to each flask resulting in a final concentration of Avicel of 1% (w/v) Unless otherwise indicated, the strains were grown and maintained on *Schizophyllum commune* minimal medium (SCMM) (Van Peer et al., 2009).

To determine the mushroom development phenotype, the *S. commune* dikaryotic strains were grown at 25°C, in a 16 h light/ 8 h dark cycle, on SCMM plates for 7 days. For the high CO_2_ experiment, the strains were grown in 5% CO_2_, continuous light.

### Genome sequencing, annotation and analysis

For all genome sequences generated in this study (Table 1), mycelium was grown from plug inoculum on top of a Poretics™ Polycarbonate Track Etched (PCTE) Membrane (GVS, Italy) on solid SCMM for 5 – 7 days at 30 °C. The resulting biomass was frozen, lyophilized and powdered. Genomic DNA isolation was performed using ChargeSwitch™ gDNA Plant Kit (Invitrogen) according to the manufacturer’s instructions and the quality and concentration were checked using agarose gel electrophoresis, NanoDrop and Qubit.

The genomes were sequenced using the Illumina platform. Plate-based DNA library preparation for Illumina sequencing was performed on the PerkinElmer Sciclone NGS robotic liquid handling system using Kapa Biosystems library preparation kit (Roche). 200 ng of sample DNA was sheared to 300 bp or 600 bp using a Covaris LE220 focused-ultrasonicator. The sheared DNA fragments were size selected by double-SPRI using TotalPure NGS beads (Omega Bio-tek) and then the selected fragments were end-repaired, A-tailed, and ligated with Illumina compatible sequencing adaptors from IDT, Inc. containing a unique molecular index barcode for each sample library. The prepared libraries were then quantified using KAPA Illumina library quantification kit (Roche) and run on a LightCycler 480 real-time PCR instrument (Roche). The quantified libraries were then multiplexed and the pool of libraries was then prepared for sequencing on the Illumina NovaSeq 6000 sequencing platform using NovaSeq XP v1 reagent kits (Illumina), S4 flow cell, following a 2×150 indexed run recipe. The raw Illumina sequence data was filtered for artifact/process contamination using the JGI QC pipeline. The genome assembly was generated using SPAdes v3.12.0 or v3.15.2 (Bankevich et al., 2012) [--phred-offset 33 --cov-cutoff auto -t 16 -m 115 -k 25,55,95 --careful --12]. The assembled genomes were annotated using the JGI Annotation pipeline (Grigoriev et al., 2014).

The genomes of the previously published strains JCM22674, Mos and FL were included for comparative purposes. The genome assembly of strain JCM22674 was obtained from NCBI GenBank BCGZ00000000.1 and was previously sequenced by RIKEN BioResource Center and RIKEN Center for Life Science Technologies through the Genome Information Upgrading Program of the National Bio-Resource Project of the MEXT, Japan. For strains Mos and FL, the genomes were de novo assembled. The raw sequencing reads were obtained from NCBI SRA accession SRR1120868 and SRR1140996, respectively. The reads were quality trimmed on both ends using BBduk to a minimum quality value of 10. The trimmed reads were assembled using SPAdes version 3.11.1 (Bankevich et al., 2012) using the ‘careful’ setting and k-mer lengths of 21, 33, 55, 77 and 99. Scaffolds shorter than 1 kb were removed. For the three strains, genes were predicted using Augustus with *S. commune*-specific parameter files that were previously generated by Braker (previously published RNA-Seq reads (Pelkmans et al., 2017) were aligned to the assembly of strain H4-8 (Marian et al., 2022) using Hisat2 (Kim et al., 2015) and Braker version 2.0.5 (Hoff et al., 2019) was subsequently run using the ‘fungus’ parameter).

Repetitive sequences in the assemblies were identified *de novo* with RepeatModeler version 2.0.3 (Flynn et al., 2020), which uses RepeatScout version 1.0.6 (Price et al., 2005), RECON version 1.08 and Tandem Repeats Finder version 4.09 (Benson, 1999). RepeatMasker version 4.1.2 (http://www.repeatmasker.org) was subsequently used to annotate the identified repetitive content, using the default Dfam database provided with RepeatMasker.

The genome conservation between each pairwise combination of assemblies was determined by aligning the assemblies using PROmer (in the software package MUMmer version 3 (Kurtz et al., 2004)) with the setting ‘mum’. The sequence similarity was calculated for each 10 kbp window. Only reference sequences longer than 10 kbp were included. The average of these windows was taken as the average similarity between the assemblies.

The predicted genes of all strains were functionally annotated. PFAM version 35 was used to predict conserved protein domains (Mistry et al., 2021) together with its corresponding gene ontology (GO) terms (Ashburner et al., 2000; Hunter et al., 2009). Signalp 4.1 (Petersen et al., 2011) and TMHMM 2.0c (Krogh et al., 2001) were used to predict secretion signals and transmembrane domains, the following criteria were applied to consider a certain protein as small, secreted protein: A) they had a secretion signal, but no transmembrane domain (except in the first 40 amino acids) and B) were shorter than 300 amino acids. Transcription factors were identified based on the presence of a PFAM domain with DNA binding properties (Park et al., 2008). CAZymes were annotated with the standalone version of the dbCAN pipeline using HMMdb version 9 (Zhang et al., 2018). The hydrophobins were identified based on the PFAM domain PF01185. A gene tree of the hydrophobins was reconstructed by aligning the sequences using MAFFT version 7.505 (Katoh and Standley, 2013) followed by gene tree reconstruction using FastTree version 2.1.11 (Price et al., 2010). The tree was manually inspected to determine the conservation of the previously identified hydrophobins of strain H4-8 (Ohm et al., 2010). The putative mating type genes were identified based on their functional annotations. The matA locus consists of homeodomain transcription factors that generally comprise a specific homeodomain (PF05920). The matB locus consists of pheromones and pheromone receptors that were identified by PFAM domains PF08015 and PF02076, respectively. Gene family conservation was analyzed using Orthofinder2 version 2.5.4 (Emms and Kelly, 2019) with an inflation parameter of 1.5. The resulting orthogroups were classified depending on their conservation.

The phylogenetic tree of species and strains was calculated by first aligning the amino acid sequences of conserved proteins identified by BUSCO using MAFFT version 7.505 (Katoh and Standley, 2013). Next, the amino acids were replaced by their corresponding codons, effectively resulting in aligned coding sequences. Phylogenetically non-informative positions were removed by Gblocks version 0.91b (Talavera et al., 2007) and the tree was calculated with FastTree version 2.1.11 (Price et al., 2010). The species *Auriculariopsis ampla* (Almási et al., 2019) and *Fistulina hepatica* (Floudas et al., 2015) were included as outgroups and the tree was rooted on *F. hepatica*.

### Cellulase enzyme activity

The *S. commune* monokaryotic strains were first pre-cultured on a Poretics™ Polycarbonate Track Etched (PCTE) Membrane (GVS, Italy) placed on top of SCMM plates for 5 days at 30°C. The mycelium of 2 petri dishes was macerated in 90 ml MwA for 1 minute at low speed in a Waring Commercial Blender. The macerate was evenly distributed to 250 ml Erlenmeyers (30 ml each) containing 70 ml MwA with 1 g Avicel, resulting in a final concentration of Avicel of 1% (w/v). Three biological replicates were grown in an Innova incubator for 7 days at 30°C shaking at 200 rpm. After 7 days 1 mL of culture was collected and centrifuged at 9391 g for 10 min. The cellulase activity was measured in the supernatant with the filter paper assay (Xiao et al., 2004). Briefly, the total cellulase activity was determined by an enzymatic reaction employing 7 mm circles of Whatman No.1 filter paper and 60 μl of supernatant. The reaction was incubated at 50°C for 72 hours. Next, 120 μL of dinitrosalicylic acid (DNS) was added to the reaction, which was then heated at 95°C for 5 minutes. Finally, 100 μl of each sample was transferred to the wells of a flat-bottom plate and absorbance was read at 540 nm using a BioTek Synergy HTX Microplate Reader. One enzyme unit (FPU) was defined as the amount of enzyme capable of liberating reducing sugar at a rate of 1 μmol min^-1^ (as determined by comparison to a glucose standard curve).

## Supplementary Data

**Table S1**. Statistics of the genome assemblies, predicted genes and functional annotations of the *S. commune* strains, *S. fasciatum*, and the outgroups *A. ampla* and *F. hepatica*.

**Table S2**. Predicted mating type genes, identified based on their functional annotation.

**Table S3**. The genes annotated in the various CAZyme families.

**Figure S1.**
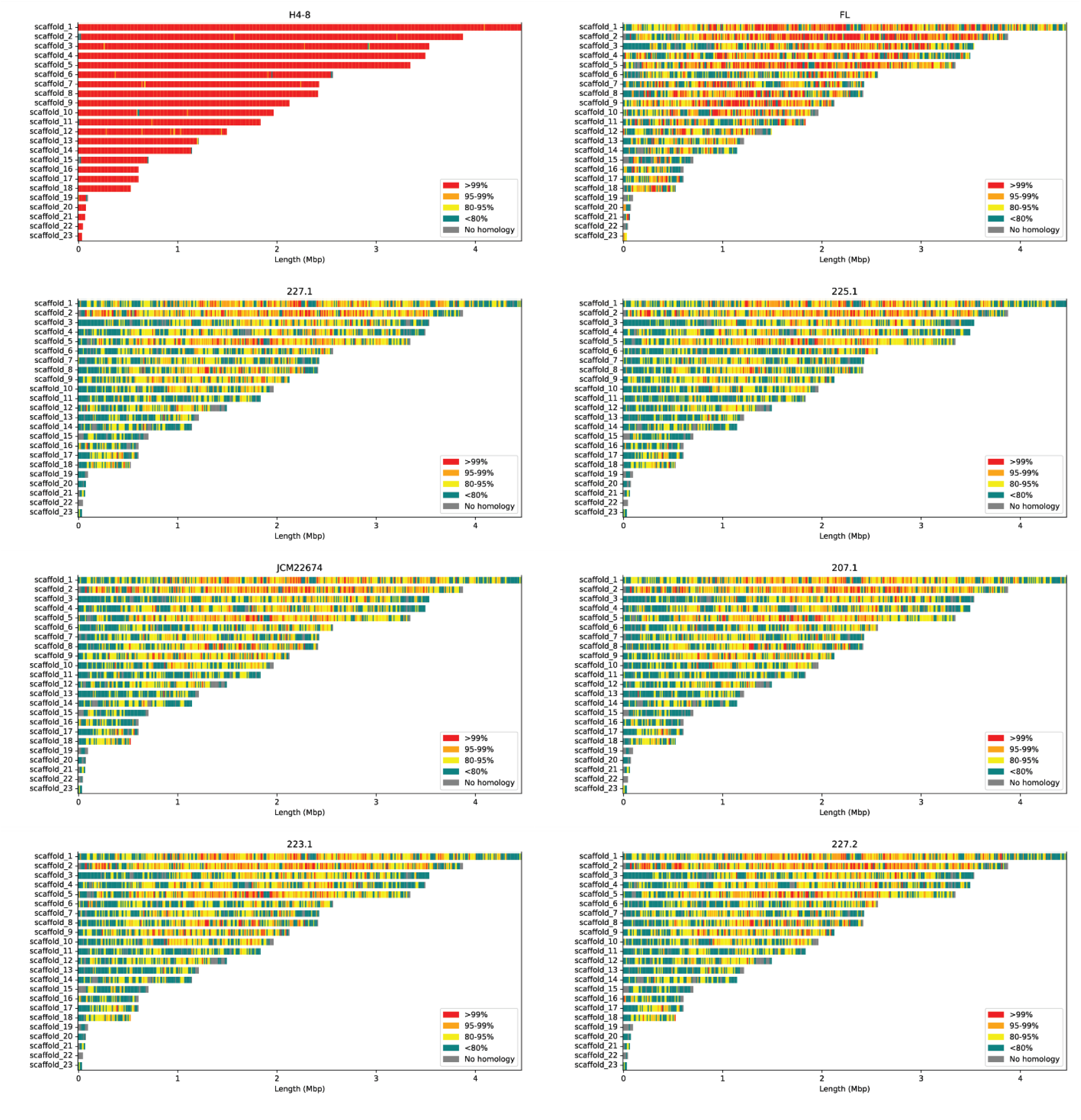
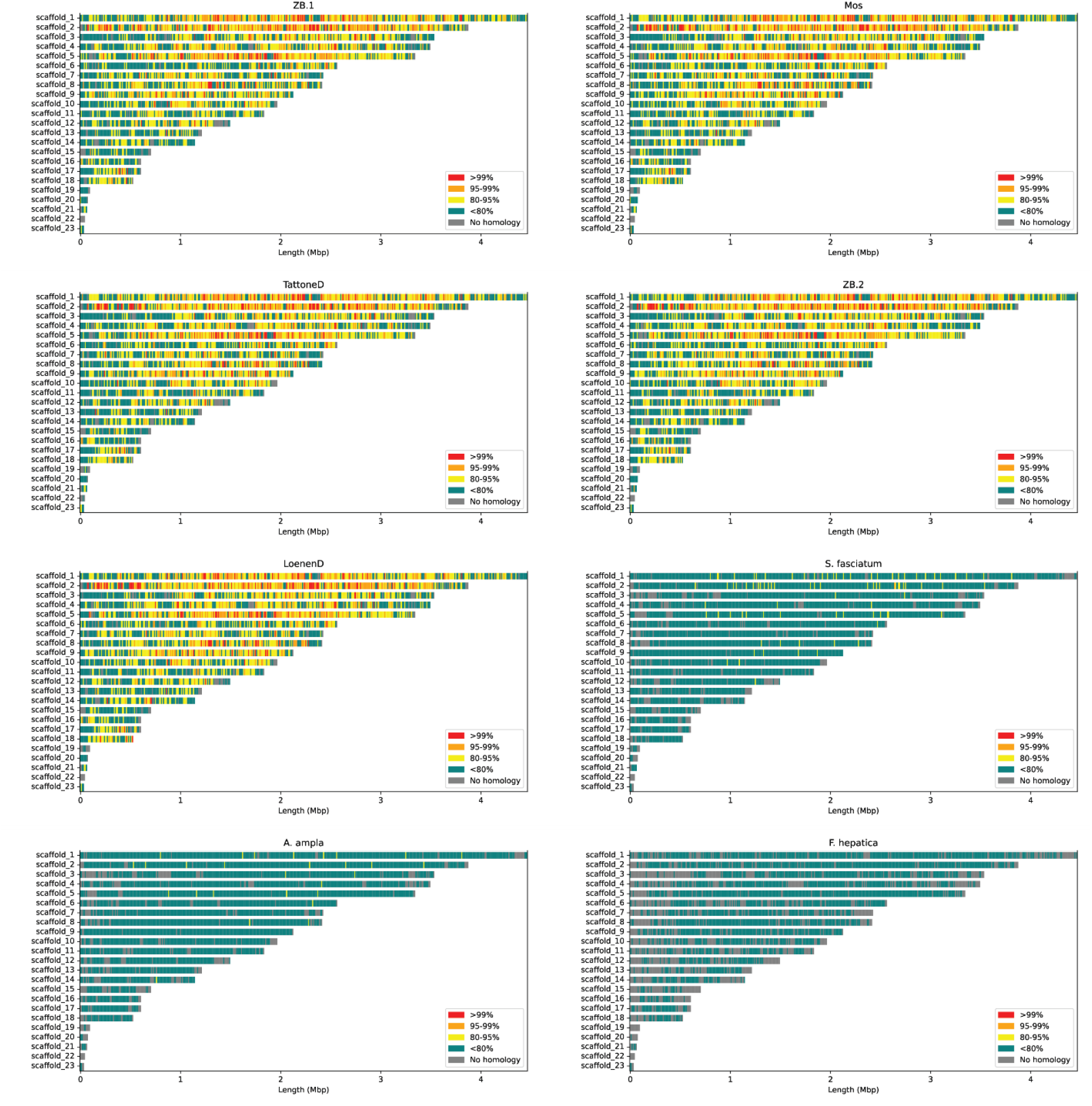
Conservation between the reference assembly of strain H4-8 and the assemblies of the other 12 *S. commune* strains, *S. fasciatum, A. ampla* and *F. hepatica*. The scaffolds of the reference assembly are visualized along the y-axis, and the scaffold lengths are indicated on the x-axis.

## Data Availability

The genome assemblies and annotations that were produced for this study are available from the JGI Fungal genome portal MycoCosm (Grigoriev et al., 2014) and have been deposited to NCBI GenBank: JAZHXD000000000 (*S. commune* 225.1), JAZHXE000000000 (*S. commune* 227.1), JAZHXF000000000 (*S. commune* ZB.1), JAZHXG000000000 (*S. commune* ZB.2), JAZHXC000000000 (*S. fasciatum* CBS 267.60 monokaryon) (remaining accessions to be provided upon publication).

## Acknowledgements

This project has received funding from the European Research Council (ERC) under the European Union’s Horizon 2020 research and innovation programme (grant agreement number 716132). The work (proposal: 10.46936/10.25585/60001043) conducted by the U.S. Department of Energy Joint Genome Institute (https://ror.org/04xm1d337), a DOE Office of Science User Facility, is supported by the Office of Science of the U.S. Department of Energy operated under Contract No. DE-AC02-05CH11231.

## References

Almási, É., Sahu, N., Krizsán, K., Bálint, B., Kovács, G.M., Kiss, B., Cseklye, J., Drula, E., Henrissat, B., Nagy, I., Chovatia, M., Adam, C., LaButti, K., Lipzen, A., Riley, R., Grigoriev, I. V., Nagy, L.G., 2019. Comparative genomics reveals unique wood-decay strategies and fruiting body development in the Schizophyllaceae. New Phytologist 224, 902–915. 10.1111/nph.16032

Ashburner, M., Ball, C.A., Blake, J.A., Botstein, D., Butler, H., Cherry, J.M., Davis, A.P., Dolinski, K., Dwight, S.S., Eppig, J.T., Harris, M.A., Hill, D.P., Issel-Tarver, L., Kasarskis, A., Lewis, S., Matese, J.C., Richardson, J.E., Ringwald, M., Rubin, G.M., Sherlock, G., 2000. Gene Ontology: tool for the unification of biology. Nat Genet 25, 25–29. 10.1038/75556

Baranova, M.A., Logacheva, M.D., Penin, A.A., Seplyarskiy, V.B., Safonova, Y.Y., Naumenko, S.A., Klepikova, A. V, Gerasimov, E.S., Bazykin, G.A., James, T.Y., Kondrashov, A.S., 2015. Extraordinary Genetic Diversity in a Wood Decay Mushroom. Mol Biol Evol 32, 2775–2783. 10.1093/molbev/msv153

Benson, G., 1999. Tandem repeats finder: a program to analyze DNA sequences. Nucleic Acids Res 27, 573–580. 10.1093/NAR/27.2.573

Boiko, S.M., 2022. Identification of novel SSR markers for predicting the geographic origin of fungus Schizophyllum commune Fr. Fungal Biol 126, 764–774. 10.1016/J.FUNBIO.2022.09.005

De Jong, J.F., Ohm, R.A., De Bekker, C., Wösten, H.A.B., Lugones, L.G., 2010. Inactivation of ku80 in the mushroom-forming fungus Schizophyllum commune increases the relative incidence of homologous recombination. FEMS Microbiol Lett 310, 91–95. 10.1111/j.1574-6968.2010.02052.x

Drula, E., Garron, M.L., Dogan, S., Lombard, V., Henrissat, B., Terrapon, N., 2022. The carbohydrate-active enzyme database: functions and literature. Nucleic Acids Res 50, D571–D577. 10.1093/NAR/GKAB1045

Emms, D.M., Kelly, S., 2019. OrthoFinder: Phylogenetic orthology inference for comparative genomics. Genome Biol 20, 238. 10.1186/s13059-019-1832-y

Floudas, D., Bentzer, J., Ahrén, D., Johansson, T., Persson, P., Tunlid, A., 2020. Uncovering the hidden diversity of litter-decomposition mechanisms in mushroom-forming fungi. The ISME Journal 2020 14:8 14, 2046–2059. 10.1038/s41396-020-0667-6

Floudas, D., Held, B.W., Riley, R., Nagy, L.G., Koehler, G., Ransdell, A.S., Younus, H., Chow, J., Chiniquy, J., Lipzen, A., Tritt, A., Sun, H., Haridas, S., LaButti, K., Ohm, R.A., Kües, U., Blanchette, R.A., Grigoriev, I. V., Minto, R.E., Hibbett, D.S., 2015. Evolution of novel wood decay mechanisms in Agaricales revealed by the genome sequences of Fistulina hepatica and Cylindrobasidium torrendii. FungalGenetics and Biology 76, 78–92. 10.1016/j.fgb.2015.02.002

Flynn, J.M., Hubley, R., Goubert, C., Rosen, J., Clark, A.G., Feschotte, C., Smit, A.F., 2020. RepeatModeler2 for automated genomic discovery of transposable element families. Proc Natl Acad Sci U S A 117, 9451–9457. 10.1073/PNAS.1921046117/SUPPL_FILE/PNAS.1921046117.SAPP.PDF

Grigoriev, I. V., Martinez, D.A., Salamov, A.A., 2006. Fungal Genomic Annotation. Applied Mycology and Biotechnology 6, 123–142. 10.1016/S1874-5334(06)80008-0

Grigoriev, I. V., Nikitin, R., Haridas, S., Kuo, A., Ohm, R., Otillar, R., Riley, R., Salamov, A., Zhao, X., Korzeniewski, F., Smirnova, T., Nordberg, H., Dubchak, I., Shabalov, I., 2014. MycoCosm portal: gearing up for 1000 fungal genomes. Nucleic Acids Res 42, D699–D704. 10.1093/nar/gkt1183

Hoff, K.J., Lomsadze, A., Borodovsky, M., Stanke, M., 2019. Whole-Genome Annotation with BRAKER. Methods Mol Biol 1962, 65. 10.1007/978-1-4939-9173-0_5

Hunter, S., Apweiler, R., Attwood, T.K., Bairoch, A., Bateman, A., Binns, D., Bork, P., Das, U., Daugherty, L., Duquenne, L., Finn, R.D., Gough, J., Haft, D., Hulo, N., Kahn, D., Kelly, E., Laugraud, A., Letunic, I., Lonsdale, D., Lopez, R., Madera, M., Maslen, J., McAnulla, C., McDowall, J., Mistry, J., Mitchell, A., Mulder, N., Natale, D., Orengo, C., Quinn, A.F., Selengut, J.D., Sigrist, C.J.A., Thimma, M., Thomas, P.D., Valentin, F., Wilson, D., Wu, C.H., Yeats, C., 2009. InterPro: the integrative protein signature database. Nucleic Acids Res 37, D211–D215. 10.1093/nar/gkn785

James, T.Y., Moncalvo, J.-M., Li, S., Vilgalys, R., 2001. Polymorphism at the ribosomal DNA spacers and its relation to breeding structure of the widespread mushroom Schizophyllum commune. Genetics 157, 149. 10.1093/GENETICS/157.1.149

James, T.Y., Porter, D., Hamrick, J.L., Vilgalys, R., 1999. Evidence for limited intercontinental gene flow in the cosmopolitan mushroom, Schizophyllum commune. Evolution (N Y) 53, 1665–1677.

Katoh, K., Standley, D.M., 2013. MAFFT Multiple Sequence Alignment Software Version 7: Improvements in Performance and Usability. Mol Biol Evol 30, 772–780. 10.1093/molbev/mst010

Kim, D., Langmead, B., Salzberg, S.L., 2015. HISAT: a fast spliced aligner with low memory requirements. Nat Methods 12, 357–360. 10.1038/nmeth.3317

Kim, D.W., Nam, J., Nguyen, H.T.K., Lee, J., Choi, Y., Choi, J., 2021. Draft Genome Sequence of the White-Rot Fungus Schizophyllum Commune IUM1114-SS01. Mycobiology 49, 86–88. 10.1080/12298093.2020.1843222

Krizsán, K., Almási, É., Merényi, Z., Sahu, N., Virágh, M., Kószó, T., Mondo, S., Kiss, B., Bálint, B., Kües, U., Barry, K., Cseklye, J., Hegedüs, B., Henrissat, B., Johnson, J., Lipzen, A., Ohm, R.A., Nagy, I., Pangilinan, J., Yan, J., Xiong, Y., Grigoriev, I. V., Hibbett, D.S., Nagy, L.G., 2019. Transcriptomic atlas of mushroom development reveals conserved genes behind complex multicellularity in fungi. Proc Natl Acad Sci U S A 116, 7409–7418. 10.1073/pnas.1817822116

Krogh, A., Larsson, B., von Heijne, G., Sonnhammer, E.L.L., 2001. Predicting transmembrane protein topology with a hidden markov model: application to complete genomes. J Mol Biol 305, 567–580. 10.1006/JMBI.2000.4315

Kurtz, S., Phillippy, A., Delcher, A.L., Smoot, M., Shumway, M., Antonescu, C., Salzberg, S.L., 2004. Versatile and open software for comparing large genomes. Genome Biol 5, R12. 10.1186/gb-2004-5-2-r12

Liu, X., Zainul Arifeen, M., Xue, Y., Liu, C., 2022. Genome-wide characterization of laccase gene family in Schizophyllum commune 20R-7-F01, isolated from deep sediment 2 km below the seafloor. Front Microbiol 13, 923451. 10.3389/FMICB.2022.923451/BIBTEX

Marian, I.M., Vonk, P.J., Valdes, I.D., Barry, K., Bostock, B., Carver, A., Daum, C., Lerner, H., Lipzen, A., Park, H., Schuller, M.B.P., Tegelaar, M., Tritt, A., Schmutz, J., Grimwood, J., Lugones, L.G., Choi, I.-G., Wösten, H.A.B., Grigoriev, I. V., Ohm, R.A., 2022. The Transcription Factor Roc1 Is a Key Regulator of Cellulose Degradation in the Wood-Decaying Mushroom Schizophyllum commune. mBio. 10.1128/MBIO.00628-22

Mistry, J., Chuguransky, S., Williams, L., Qureshi, M., Salazar, G.A., Sonnhammer, E.L.L., Tosatto, S.C.E., Paladin, L., Raj, S., Richardson, L.J., Finn, R.D., Bateman, A., 2021. Pfam: The protein families database in 2021. Nucleic Acids Res 49, D412–D419. 10.1093/NAR/GKAA913

Niederpruem, D.J., 1963. Role of carbon dioxide in the control of fruiting of Schizophyllum commune. J Bacteriol 85, 1300–1308. 10.1128/JB.85.6.1300-1308.1963

Ohm, R.A., Aerts, D., Wösten, H.A.B., Lugones, L.G., 2013. The blue light receptor complex WC-1/2 of Schizophyllum commune is involved in mushroom formation and protection against phototoxicity. Environ Microbiol 15, 943–955. 10.1111/j.1462-2920.2012.02878.x

Ohm, R.A., de Jong, J.F., de Bekker, C., Wösten, H.A.B., Lugones, L.G., 2011. Transcription factor genes of Schizophyllum commune involved in regulation of mushroom formation. Mol Microbiol 81, 1433–1445. 10.1111/j.1365-2958.2011.07776.x

Ohm, R.A., De Jong, J.F., Lugones, L.G., Aerts, A., Kothe, E., Stajich, J.E., De Vries, R.P., Record, E., Levasseur, A., Baker, S.E., Bartholomew, K.A., Coutinho, P.M., Erdmann, S., Fowler, T.J., Gathman, A.C., Lombard, V., Henrissat, B., Knabe, N., Kües, U., Lilly, W.W., Lindquist, E., Lucas, S., Magnuson, J.K., Piumi, F., Raudaskoski, M., Salamov, A., Schmutz, J., Schwarze, F.W.M.R., vanKuyk, P.A., Horton, J.S., Grigoriev, I. V, Wösten, H.A.B., 2010. Genome sequence of the model mushroom Schizophyllum commune. Nat Biotechnol 28, 957–963.

Ohm, R.A., Riley, R., Salamov, A., Min, B., Choi, I.-G., Grigoriev, I. V., 2014. Genomics of wood-degrading fungi. Fungal Genetics and Biology 72, 82–90. 10.1016/J.FGB.2014.05.001

Park, Jongsun, Park, Jaejin, Jang, S., Kim, Seryun, Kong, S., Choi, J., Ahn, K., Kim, J., Lee, S., Kim, Sunggon, Park, B., Jung, K., Kim, Soonok, Kang, S., Lee, Y.-H., 2008. FTFD: an informatics pipeline supporting phylogenomic analysis of fungal transcription factors. Bioinformatics 24, 1024–1025. 10.1093/bioinformatics/btn058

Pelkmans, J.F., Patil, M.B., Gehrmann, T., Reinders, M.J.T., Wösten, H.A.B., Lugones, L.G., 2017. Transcription factors of Schizophyllum commune involved in mushroom formation and modulation of vegetative growth. Sci Rep 7, 1–11. 10.1038/s41598-017-00483-3

Petersen, T.N., Brunak, S., von Heijne, G., Nielsen, H., 2011. SignalP 4.0: discriminating signal peptides from transmembrane regions. Nat Methods 8, 785–786. 10.1038/nmeth.1701

Price, A.L., Jones, N.C., Pevzner, P.A., 2005. De novo identification of repeat families in large genomes. Bioinformatics 21, i351–i358. 10.1093/bioinformatics/bti1018

Price, M.N., Dehal, P.S., Arkin, A.P., 2010. FastTree 2 – Approximately Maximum-Likelihood Trees for Large Alignments. PLoS One 5, e9490. 10.1371/journal.pone.0009490

Raudaskoski, M., Viitanen, H., 1982. Effect of aeration and light on fruit body induction in Schizophyllum commune. Transactions of the British Mycological Society 78, 89–96. 10.1016/S0007-1536(82)80080-6

Riley, R., Salamov, A.A., Brown, D.W., Nagy, L.G., Floudas, D., Held, B.W., Levasseur, A., Lombard, V., Morin, E., Otillar, R., Lindquist, E.A., Sun, H., LaButti, K.M., Schmutz, J., Jabbour, D., Luo, H., Baker, S.E., Pisabarro, A.G., Walton, J.D., Blanchette, R.A., Henrissat, B., Martin, F., Cullen, D., Hibbett, D.S., Grigoriev, I. V, 2014. Extensive sampling of basidiomycete genomes demonstrates inadequacy of the white-rot/brown-rot paradigm for wood decay fungi. Proceedings of the National Academy of Sciences 111, 9923–9928. 10.1073/pnas.1400592111

Seplyarskiy, V.B., Logacheva, M.D., Penin, A.A., Baranova, M.A., Leushkin, E. V., Demidenko, N. V., Klepikova, A. V., Kondrashov, F.A., Kondrashov, A.S., James, T.Y., 2014. Crossing-Over in a Hypervariable Species Preferentially Occurs in Regions of High Local Similarity. Mol Biol Evol 31, 3016–3025. 10.1093/MOLBEV/MSU242

Talavera, G., Castresana, J., Kjer, K., Page, R., Sullivan, J., 2007. Improvement of Phylogenies after Removing Divergent and Ambiguously Aligned Blocks from Protein Sequence Alignments. Syst Biol 56, 564–577. 10.1080/10635150701472164

Van Peer, A.F., De Bekker, C., Vinck, A., Wösten, H.A.B., Lugones, L.G., 2009. Phleomycin increases transformation efficiency and promotes single integrations in Schizophyllum commune. Appl Environ Microbiol 75, 1243–1247. 10.1128/AEM.02162-08

Vonk, P.J., Escobar, N., Wösten, H.A.B., Lugones, L.G., Ohm, R.A., 2019. High-throughput targeted gene deletion in the model mushroom Schizophyllum commune using pre-assembled Cas9 ribonucleoproteins. Sci Rep 9, 7632. 10.1038/s41598-019-44133-2

Vonk, P.J., Ohm, R.A., 2021. H3K4me2 ChIP-Seq reveals the epigenetic landscape during mushroom formation and novel developmental regulators of Schizophyllum commune. Sci Rep 11, 8178. 10.1038/s41598-021-87635-8

Vonk, P.J., Ohm, R.A., 2018. The role of homeodomain transcription factors in fungal development. Fungal Biol Rev 32, 219–230. 10.1016/J.FBR.2018.04.002

Whitaker, D.R., 1951. Studies in the biochemistry of cellulolytic microorganisms: I. carbon balances of wood-rotting fungi in surface culture. 10.1139/b51-016 29, 159–175. 10.1139/B51-016

Wösten, H.A.B., 2001. Hydrophobins: Multipurpose Proteins. Annu Rev Microbiol 55, 625–646. 10.1146/annurev.micro.55.1.625

Wu, B., Xu, Z., Knudson, A., Carlson, A., Chen, N., Kovaka, S., LaButti, K., Lipzen, A., Pennachio, C., Riley, R., Schakwitz, W., Umezawa, K., Ohm, R.A., Grigoriev, I. V., Nagy, L.G., Gibbons, J., Hibbett, D., 2018. Genomics and Development of Lentinus tigrinus: A White-Rot Wood-Decaying Mushroom with Dimorphic Fruiting Bodies. Genome Biol Evol 10, 3250–3261. 10.1093/GBE/EVY246

Xiao, Z., Storms, R., Tsang, A., 2004. Microplate-based filter paper assay to measure total cellulase activity. Biotechnol Bioeng 88, 832–837. 10.1002/bit.20286

Zhang, H., Yohe, T., Huang, L., Entwistle, S., Wu, P., Yang, Z., Busk, P.K., Xu, Y., Yin, Y., 2018. DbCAN2: A meta server for automated carbohydrate-active enzyme annotation. Nucleic Acids Res 46, W95–W101. 10.1093/nar/gky418

